# No evidence that post-training D2R dopaminergic drug administration affects fear generalization in male rats

**DOI:** 10.1101/2022.02.02.478781

**Authors:** Natalie Schroyens, Laura Vercammen, Burcu Özcan, Victoria Aurora Ossorio Salazar, Jonas Zaman, Dimitri De Bundel, Tom Beckers, Laura Luyten

**Affiliations:** KU Leuven, Centre for the Psychology of Learning and Experimental Psychopathology, Faculty of Psychology and Educational Sciences, Tiensestraat 102 box 3712, 3000 Leuven, Belgium; KU Leuven, Leuven Brain Institute, O&N V Herestraat 49 box 1020, 3000 Leuven, Belgium; KU Leuven, Laboratory of Biological Psychology, Faculty of Psychology and Educational Sciences, Tiensestraat 102 box 3714, 3000 Leuven, Belgium; KU Leuven, Health Psychology, Tiensestraat 102 box 3726, 3000 Leuven, Belgium; Research Group Experimental Pharmacology, Department of Pharmaceutical Sciences, Center for Neurosciences, Vrije Universiteit Brussel, 1090 Brussel, Belgium

**Keywords:** differential cued fear conditioning, generalization, rats, dopamine, quinpirole, raclopride

## Abstract

The neurotransmitter dopamine plays an important role in the processing of emotional memories, and prior research suggests that dopaminergic manipulations immediately after fear learning can affect the retention and generalization of acquired fear. The current study focuses specifically on the role of dopamine D2 receptors (D2Rs) in adult, male Wistar rats. In a series of five experiments, D2R (ant)agonists were injected systemically immediately after differential cued fear conditioning (CS+ followed by shock, CS− without shock). All five experiments involved the administration of the D2R agonist quinpirole at different doses versus saline (n = 12, 16 or 44 rats/group). Additionally, one of the studies administered the D2R antagonist raclopride (n = 12). One day later, freezing during the CS+ and CS− was assessed. We found no indications for an effect of quinpirole or raclopride on fear generalization during this drug-free test. Importantly, and contradicting prior research in mice, the evidence for the absence of an effect of quinpirole (1 mg/kg) on fear generalization was substantial according to Bayesian analyses and was observed in a highly powered experiment (N = 87). We did find acute behavioral effects in line with the literature, for both quinpirole and raclopride in a locomotor activity test.

## 1. Introduction

The neurotransmitter dopamine has been shown to play a role in the processing of emotional memories^1^. For instance, genetic and pharmacological studies suggest that activation of dopaminergic D1 and D2 receptors immediately after Pavlovian conditioning or inhibitory avoidance training is required for subsequent expression of acquired fear^2–6^. In addition, dopamine has been found to modulate long-term potentiation^7–11^, a form of synaptic plasticity that is assumed to be essential for memory formation^12^. The focus of the present paper is on the role of dopaminergic signaling after fear conditioning with two cues, differentially predicting danger (CS+) and the absence of danger (CS−), in subsequent fear generalization from the CS+ to the CS−.

Although most studies regarding the influence of dopamine on fear memory processing have focused on its effects on memory retention specifically, more recent studies have also investigated its role in generalization of fear towards safe or novel stimuli after contextual, cued, or differential cued fear conditioning^13–17^. Investigating if and how dopaminergic manipulations during or after learning affect the retention and spreading of fear towards safe situations is highly relevant to understand the neurobiology of anxiety, given that fear generalization is often found to be elevated in patients with anxiety disorders^18,19^.

Prior research indicated that knock-out mice lacking hippocampal dopamine D1 receptors (D1Rs) exhibit equal levels of freezing during exposure to a novel context as to a context that was previously paired with shock^16^. In addition, using differential cued fear conditioning procedures in mice, it has been shown that fear generalization from the CS+ to CS− can be countered by optogenetic stimulation of dopaminergic neurons during conditioning^14^ or by systemic injection of a D2R agonist immediately after conditioning^13^. The latter study, by De Bundel and colleagues, found the dopaminergic effect to be bidirectional, with post-learning injection or intra-amygdala infusion of a D2R antagonist increasing the generalization from the CS+ to the CS− relative to the saline groups in these experiments. A PET imaging study in humans^20^ found evidence consistent with these findings in mice. Likewise, recent rat studies (often with more elaborate behavioral procedures that also included reward cues, or using compound or contextual stimuli, rather than simple differential learning with a tone CS+ and CS−^21–24^) have provided accumulating evidence for the involvement of dopamine signaling in the discrimination between danger and the absence of danger. The pharmacological and recording techniques that were used in these rat studies did, however, not allow for conclusions regarding the specific involvement of D2 receptors.

The abovementioned neurobiological studies have proposed a range of different mechanisms that may underlie the effect of dopamine on the spreading of fear towards safe situations. For example, it has been suggested that the involvement of dopamine in the encoding of novel perceptual information may lie at the basis of its effects on fear generalization^16^. Alternatively, it has been proposed that the generalized fear observed in knockout mice with impaired dopaminergic signaling can be attributed to inadequate learning of CS− US associations due to aberrant salience detection^17^ and the erratic assignment of negative valence to both the CS+ and CS− during learning^14^. Yet another possible way in which dopamine may limit fear generalization from stimuli that predict danger to those that predict safety is via enhancement of the stabilization or ‘consolidation’ of the acquired fear memory^13^. Related to this idea, it has been suggested that ensuring adequate stabilization of the fear memory trace may be required for the establishment of a sufficiently precise memory to prevent generalization to non-threatening situations^25^. Overall, dopamine’s effects on fear memory generalization have thus been attributed to several aspects of learning (i.e., encoding of perceptual information, association formation, differential learning), at different points in time (i.e., during versus shortly after learning), and via various hypothetical mechanisms (i.e., by coding salience or valence, or by enhancing memory stabilization).

Our ultimate goal was to study, in rats rather than mice, to what extent dopamine D2 signaling after differential fear learning plays a role in subsequent fear generalization and, if so, which aspect(s) of learning (perceptual, associative, and/or differential) would be involved. To this aim, we first embarked on a conceptual replication of published findings in mice by De Bundel and colleagues^13^, in which it was shown that systemic injection of a D2R agonist or antagonist immediately after differential cued fear conditioning can attenuate or augment subsequent fear generalization, respectively (i.e., decrease or increase fear responding to a safe cue in a new context). One advantage of administering dopaminergic drugs after learning rather than before is that it allows to distinguish between the importance of dopaminergic signaling during versus after learning. It may thus provide useful insight into the role of dopamine during the memory stabilization phase specifically.

While most prior evidence in non-human animals regarding the role of dopamine D2Rs in attenuating or augmenting fear generalization has been obtained in mice, the translation of those findings to rats remains unclear. Despite the many similarities between both species, rats and mice are characterized by important differences in functional brain anatomy and physiology, including neurochemistry^26^. Rats are an important model organism for behavioral neuroscience, in particular in the study of fear learning and interventions to influence fear memory specificity. Moreover, establishing the generalizability of the findings in mice to rats may also inform us about their potential relevance for other organisms, including humans. Exactly copying behavioral procedures from mice to rats is, however, rarely advisable, and we slightly adapted the fear learning tasks from De Bundel et al.^13^ in order to make them more appropriate for rats and less sensitive to order and context effects.

We describe a series of five experiments (Exp. 1-5) that investigated the behavioral effects of post-training dopaminergic drug injection on subsequent fear generalization in rats. These experiments were preceded by a series of protocol optimization studies (reported as Exp. A-C) that provided insight into the effects of shock intensity and testing order on fear responding to the CS+ and CS−. In a last experiment (Exp. 6), the acute effects on locomotor activity of the drugs used throughout our experiments was evaluated (i.e., 0.3 mg/kg raclopride and 0.05, 1, and 5 mg/kg quinpirole). All details regarding Experiments A-B-C and Experiment 6 can be found in the **Supplement**.

## 2. Materials and Methods

### 2.1. Protocol registration

All study protocols and statistical analyses were preregistered on the Open Science Framework (OSF) at https://osf.io/kzcyt.

### 2.2. Availability of data and materials

All datasets generated and analyzed during the current study, the analysis scripts, and results of all preregistered statistical analyses are also available on OSF (https://osf.io/kzcyt). This study is reported in accordance with ARRIVE guidelines (https://arriveguidelines.org).

### 2.3. Subjects

Male Wistar rats (Janvier Labs, Le Genest-Saint-Isle, France) were housed in groups of 3 or 4 rats per cage on a 12 h/12 h day-night cycle (lights on at 8 am). Cage enrichment, in the form of a tunnel hanging from the top grid, was provided and food and water were available ad libitum. Experiments were carried out between 9 am and 5 pm. All experiments were approved by the KU Leuven animal ethics committee, and conducted in accordance with the Belgian Royal Decree of 29/05/2013 and European Directive 2010/63/EU.

### 2.4. Drug administration

Depending on the experiment (see **Table 1**), (-)-quinpirole HCl (QUIN; 0.05 mg/kg, 1 mg/kg or 5 mg/kg; Tocris), raclopride HCl (RACLO; 0.3 mg/kg; Santa Cruz Biotechnology), or saline was administered immediately after conditioning. Both drugs were dissolved in saline (0.9%, w/v) and injected intraperitoneally at a volume of 1 ml/kg. Drug doses were based on the study in mice by De Bundel and colleagues^13^. Similar doses have previously been used in rats^27,28^.

**Table 1.**
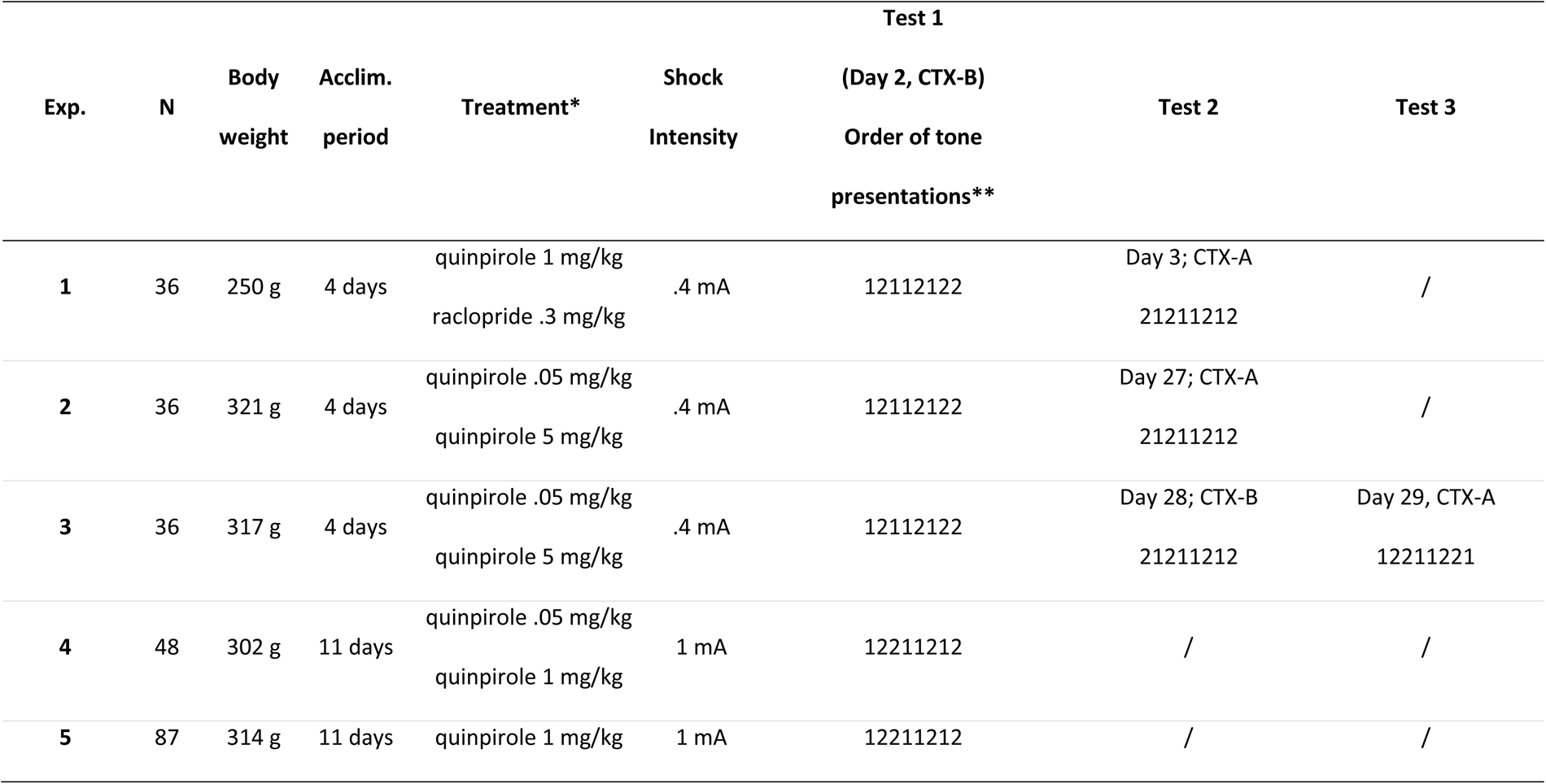
Study characteristics of Experiments 1-5. All experiments were preregistered on OSF at https://osf.io/kzcyt/registrations. Sample size (N) and average body weight one day before training are shown. Acclim. period = interval between the rats’ arrival in our animal facility and the start of handling (two consecutive days of handling preceded the fear conditioning protocol). Rats underwent differential cued fear conditioning in context A (CTX-A), after which they were injected with saline or a dopaminergic drug (quinpirole or raclopride). During subsequent drug-free sessions, they were tested in CTX-A or context B (CTX-B).*All experiments contained a saline control group. **\*\***The order of the tones at test was counterbalanced (1 = CS+, 2 = CS− or 1 = CS−, 2 = CS+).

### 2.5. Differential cued fear conditioning

Four Med Associates test chambers were used to train and test the animals. Details can be found in the **Supplement**. Fear conditioning in context A (CTX-A) consisted of presentations of a danger-signaling tone (5 x CS+, always co-terminating with a foot shock (US)) and a safe tone (5 x CS−, never paired with the shock) in a semi-random order (see **Table 1** and **Fig. 1**). Next, rats received an intraperitoneal injection of saline, quinpirole (0.05, 1 or 5 mg/kg) or raclopride (0.3 mg/kg) at a volume of 1 ml/kg. Twenty-four hours after training, rats underwent a drug-free test in a novel context (CTX-B), during which the CS+ and CS− were repeatedly presented (4 times each) in a semi-random order. Depending on the experiment, additional drug-free test sessions were performed, in either CTX-A or CTX-B.

**Fig. 1.**
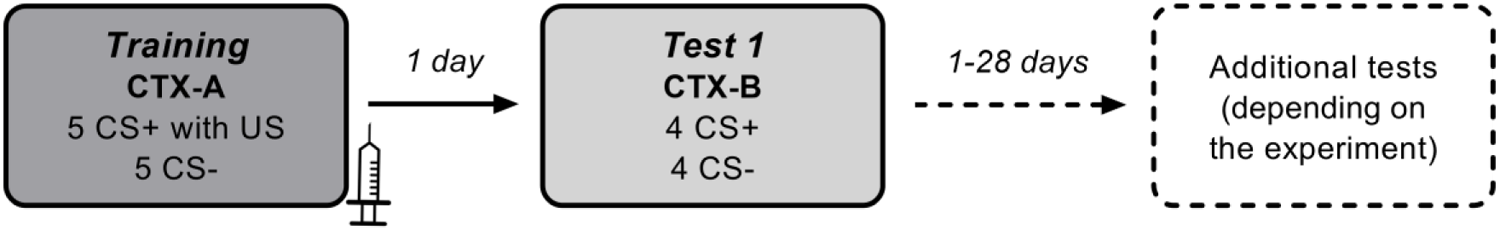
Experimental procedure. Immediately after differential cued fear conditioning in context A (CTX-A), rats were systemically injected with saline, quinpirole, or raclopride (see **Table 1** for details). Twenty-four hours later, during a drug-free test, fear responding to the CS+ and CS− was assessed in a novel context B (CTX-B, Test 1) using percentage of time spent freezing during the tones as an outcome measure, to evaluate the effects of the dopaminergic drug on subsequent fear generalization. Depending on the experiment, additional behavioral tests took place at least one day after Test 1.

### 2.6. Statistical analyses

Percentage freezing during tones was scored by an experienced observer, blinded to treatment condition, tone frequency (5 or 10 kHz) and CS type (CS− or CS+). Average % freezing during CS+ and CS− was calculated for each rat and for each test session. According to a preregistered exclusion criterion, rats showing less than 10% freezing on average during CS+ presentations at Test 1 were excluded from the analysis, as these low freezing scores may indicate a lack of acquisition of the CS− US association. Analyses were performed with and without those rats. R was used for frequentist and Bayesian analyses. For Bayesian analyses, we used a default Cauchy prior on the standardized effect size with a scale of 0.707 in the BayesFactor package. Bayes factors indicate how likely the obtained data are under H_A_, relative to H_0_ (i.e., BF_10_).

Primary analyses for Experiments 1-5 (secondary analyses are described in the **Supplement**) included mixed ANOVAs with between-subjects factor Treatment (saline or drug) and within-subjects factor CS type (CS− or CS+) to investigate whether treatment affected fear responding to both tones during tests. Mixed ANOVAs were performed for each drug (versus saline) separately. The Bayes factor (BF) quantifying the evidence for including the Treatment x CS type interaction was calculated by dividing the BF for the model with interaction and main effects by the BF for the model with only main effects, (see doi 10.31234/osf.io/spreb; referred to as ‘BFinclusion across matched models’). One-sided paired t-tests assessed whether freezing during CS+ was higher compared to CS− for each treatment condition.

Primary statistical analyses for Experiment 5 were the same as for Exp. 1-4, but in contrast with the previous experiments, this study used a sequential design, for which the total sample size depended on the value of the Bayes factor for the CS type x Treatment interaction observed during interim assessments (see **Table 2**). The first assessment took place after testing a total of 40 rats (batch 1; n = 20 rats/treatment condition) and the second assessment after testing a total of 64 rats (batch 2; 24 additional rats; n = 12 rats/treatment condition). The maximum sample size (N = 88) was determined based on frequentist power calculations using the results obtained in Experiment 4 (CS Type x Treatment interaction; power = 92%, alpha = .05, partial eta squared = .13).

**Table 2.** Overview of the sequential design used for Experiment 5. The reported power was calculated based on the effect size for the CS Type x Treatment interaction observed in Experiment 4. Interim assessments – performed after collecting data for batch 1 and 2 – included the computation of the Bayes factor for the CS type x Treatment interaction (testing stopped if the obtained BF was larger than 10 or smaller than 1/10 at interim assessment). Note that one saline rat was excluded from all analyses because it showed no freezing during any of the sessions.

| Batch | N | N<sub>total</sub> | Obtained power |
| --- | --- | --- | --- |
|  |  | (all batches combined) | (alpha = .05) |
| 1 | 40 | 40 | .60 |
| 2 | 24 | 64 | .80 |
| 3 | 24 | 88 | .92 |

## 3. Results

All details regarding the behavioral protocol optimization studies (Experiments A-B-C) that preceded Experiments 1-5 can be found in the **Supplement**.

### 3.1. Experiment 1: No evidence for an effect of post-training injection of raclopride (0.3 mg/kg) or quinpirole (1 mg/kg) on subsequent fear generalization

Based on the results of De Bundel and colleagues in mice^13^, we hypothesized that 0.3 mg/kg raclopride or 1 mg/kg quinpirole would increase or decrease fear generalization at test, respectively.

Primary analyses provided no evidence for Treatment (dopaminergic drug versus saline) x CS type (CS− versus CS+) interactions at Test 1 (RACLO: F(1, 22) = .04, p = .847, η2p < .01, BF = .38; QUIN1: F(1, 22) = .20, p = .662, η2p = .01, BF = .38). One-sided paired t-tests indicated higher freezing during CS+ than CS− presentations at test in saline control rats (t(11) = -2.42, p = .017, d = -.64, BF = 4.35), which, together with the considerable freezing levels during the CS−, suggested some differentiation, but also partial generalization at the group level. This intermediate level of generalization theoretically allowed for its augmentation or attenuation in the treatment groups. One-sided paired t-tests further indicated higher freezing during CS+ than CS− presentations at test in raclopride rats (t(11) = -2.6, p = .012, d = -.67, BF = 5.66), but not in quinpirole rats (t(11) = -1.38, p = .098, d = -.33, BF = 1.09), both in contrast with our hypotheses.

Although freezing during CS+ presentations at Test 1 was on average lower in quinpirole rats than in saline control rats (see **Fig. 2a**), two-sided t-tests (preregistered secondary analysis, see OSF) suggested no evidence for differences in freezing during CS+ presentations between treatment conditions (i.e., QUIN1 versus SAL or RACLO versus SAL). In addition, there was no evidence for differences in baseline freezing (i.e., before onset of the first tone, see OSF) or the proportion of generalizers between treatment conditions.

**Fig. 2.**
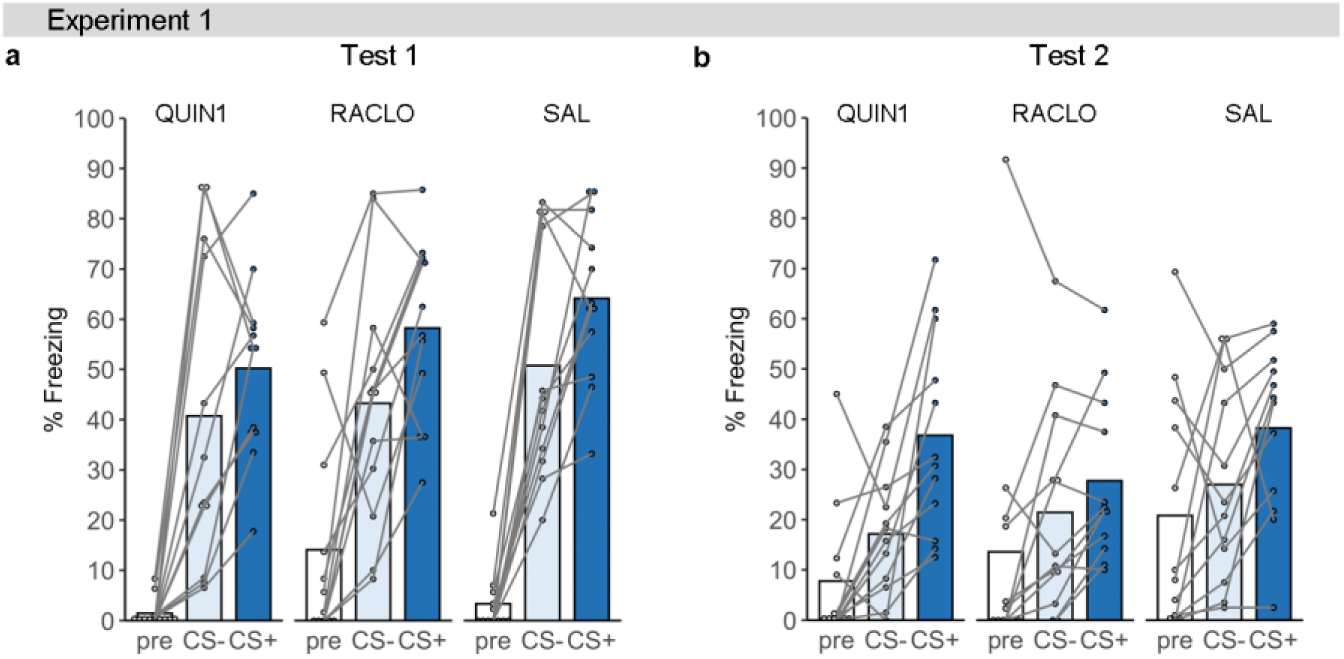
Experiment 1. Average % freezing per treatment condition during Test 1 (Day 2; CTX-B; panel a) and Test 2 (Day 3; CTX-A; panel b), shown for the 180-s baseline period (pre), CS− and CS+. QUIN1 = quinpirole (1 mg/kg), RACLO = raclopride (0.3 mg/kg), SAL = saline.

During Test 2, which took place in the training context on Day 3 (**Fig. 2b**), there was again no evidence for Treatment x CS type interactions (RACLO: F(1, 22) = .67, p = .421, η2p = .03, BF = .45; QUIN1: F(1, 22) = 1.25, p = .276, η2p = .05, BF = .63) and one-sided paired t-tests suggested higher freezing during CS+ than CS− presentations at test in saline control rats (t(11) = -2.09, p = .03, d = -.60, BF = 2.75), raclopride rats (t(11) = -2.31, p = .021, d = -.29, BF = 3.75), and in quinpirole rats (t(11) = – 3.75, p = .002, d = -1.14, BF = 30.08). Secondary analyses suggested no evidence for differences in freezing during CS+ presentations between treatment conditions (i.e., QUIN1 versus SAL or RACLO versus SAL). In addition, there was no evidence for differences in baseline freezing (i.e., before onset of the first tone), average freezing during CS+ presentations, or the proportion of generalizers between treatment conditions.

### 3.2. Experiments 2 & 3: No evidence for an effect of post-training injection of quinpirole (0.05 or 5 mg/kg) on subsequent fear generalization

Locomotor activity studies in rats suggest that quinpirole differentially affects presynaptic versus postsynaptic D2 receptors depending on the applied dose^29–32^. Experiments 2 and 3 aimed to assess how different doses of quinpirole influence fear generalization. We hypothesized that a low dose of quinpirole (0.05 mg/kg) would increase fear generalization (due to the assumed activation of presynaptic autoreceptors), whereas a higher dose of 5 mg/kg would decrease fear generalization (due to the assumed activation of postsynaptic D2Rs). Experiments 2 and 3 adopted the exact same behavioral procedures and drug doses (apart from additional tests 2 and 3), and are therefore presented together.

In Experiment 2 (**Fig. 3a-b**), three rats (1 SAL and 2 QUIN5 rats) were excluded due to the preregistered exclusion criterion of showing less than 10% freezing on average during CS+ presentations at Test 1, so all preregistered analyses were performed with and without those three rats. Statistical results are shown here for the sample in which the exclusion criterion was applied, and results for both samples (i.e., with and without exclusions) are provided only when both analyses reached different overall conclusions. All results can be found at https://osf.io/kzcyt.

**Fig. 3.**
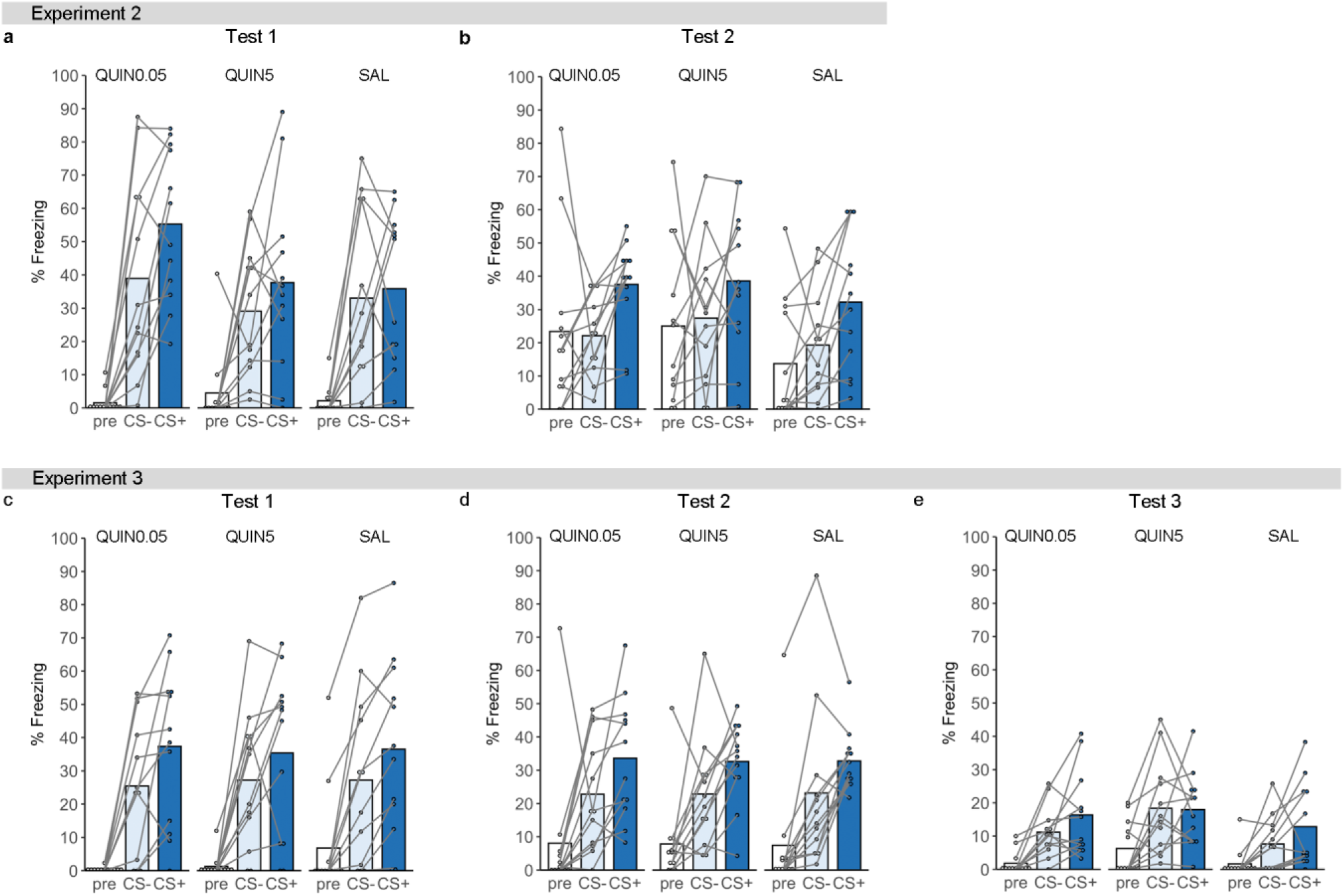
Experiments 2 & 3. Average % freezing per treatment condition during Test 1 (Day 2; CTX-B; panel a) and Test 2 (Day 27; CTX-A; panel b) of Experiment 2, and during Test 1 (Day 2; CTX-B; panel c), Test 2 (Day 28; CTX-B; panel d), and Test 3 (Day 29; CTX-A; panel e) of Experiment 3, shown for the 180-s baseline period (pre), CS− and CS+. QUIN0.05 = quinpirole (0.05 mg/kg), QUIN5 = quinpirole (5 mg/kg), SAL = saline.

Primary analyses suggested no evidence for Treatment (quinpirole versus saline) x CS type (CS− versus CS+) interactions (QUIN (0.05 mg/kg): F(1, 21) = 2.01, p = .171, η2p = .09, BF = .89; QUIN (5 mg/kg): F(1, 19) = .53, p = .477, η2p = .03, BF = .48) at Test 1. Contrary to Experiment 1, in which the same behavioral procedures were used for training and testing as here, there was no statistically significant evidence for more freezing during CS+ compared to CS− presentations at test in saline control rats (t(10) = -.39, p = .353, d = -.12, BF = .40), suggesting more complete generalization in the control group of this experiment. A similar pattern of responding was observed in rats treated with the high dose of quinpirole (t(9) = -1.34, p = .106, d = -.51, BF = 1.1). In contrast, rats that received the low quinpirole dose did show significantly more freezing during the CS+ than during the CS− (t(11) = – 2.67, p = .011, d = -.59, BF = 6.25). Neither of the quinpirole doses thus induced the hypothesized effects on fear generalization. If anything, the low quinpirole dose (0.05 mg/kg) seemed to reduce, rather than augment, fear generalization, although this pattern did not reach statistical significance.

Average freezing during CS+ presentations at Test 1 was numerically higher in the ‘low quinpirole’ condition than in saline control rats (see **Fig. 3a**), but this between-group difference did not receive statistical support when excluding the saline rat showing on average less than 10% freezing during CS+ presentations (t(21) = 1.79, p = .088, d = .75, BF = 1.16). When including all rats, the effect was statistically significant, although the obtained evidence was indifferent according to the Bayesian analysis (t(22) = 2.1, p = .047, d = .86, BF = 1.72). Furthermore, no between-group differences were found in baseline freezing (i.e., before onset of the first tone) or the proportion of generalizers.

During Test 2, which took place in the training context on Day 27 (**Fig. 3b**), there was again no evidence for Treatment x CS type interactions (QUIN (0.05 mg/kg): F(1, 21) = .06, p = .801, η2p < .01, BF = .37; QUIN (5 mg/kg): F(1, 19) < .01, p = .945, η2p < .01, BF = .38). Furthermore, both QUIN0.05 and saline rats now showed significantly higher freezing during CS+ than CS− (t(10) = -2.96, p = .007, d = -.77, BF = 8.98; t(11) = -3.85, p = .001, d = -1.20, BF = 34.85, respectively), whereas QUIN5 rats did not show statistically significant differential responding (t(9) = -1.61, p = .071, d = -.63, BF = 1.51). In addition, secondary analyses suggested no evidence for differences in baseline freezing (i.e., before onset of the first tone), average freezing during CS+ presentations, or the proportion of generalizers between treatment conditions.

In Experiment 3 (**Fig. 3c-e**), eight rats (2 SAL, 2 QUIN0.05, and 4 QUIN5 rats) were excluded due to the preregistered exclusion criterion of showing less than 10% freezing on average during CS+ presentations at Test 1, so all preregistered analyses were performed with (N = 36) and without (N = 28) those eight rats.

Primary analyses suggested no evidence for Treatment (dopaminergic drug versus saline) x CS type (CS− versus CS+) interactions (QUIN (0.05 mg/kg): F(1, 18) = .55, p = .468, η2p = .03, BF = .47; QUIN (5 mg/kg): F(1, 16) = .88, p = .362, η2p = .05, BF = .54) at Test 1 (**Fig. 3c**). One-sided paired t-tests suggested higher freezing during CS+ than CS− presentations at test in all treatment conditions (saline control rats: (t(9) = -3.64, p = .003, d = -.42, BF = 20.35; QUIN (0.05 mg/kg): t(9) = -2.82, p = .01, d = – .76, BF = 7.03; QUIN (5 mg/kg): t(7) = -3.52, p = .005, d = -.98, BF = 13.27). However, the difference in fear responding between CS+ and CS− was smaller (and non-significant) when including all QUIN5 rats (t(22) = .08, p = .933, d = .03, BF = .37), probably due to the fact that two rats showed no freezing during any of the tones, and thus, no differential responding.

During Test 2 (**Fig. 3d**), which took place in the testing context (CTX-B) on Day 28, there was again no evidence for Treatment x CS type interactions (QUIN0.05: F(1, 18) = .57, p = .458, η2p = .03, BF = .47; QUIN5: F(1, 16) = .03, p = .861, η2p < .01, BF = .39). Saline control rats (t(9) = -1.25, p = .121, d = -.24, BF = .99) and QUIN5 rats (t(7) = -1.37, p = .107, d = -.77, BF = 1.18) did no longer show significantly more freezing to the CS+ than to the CS−, whereas QUIN0.05 rats still did (t(9) = -2.76, p = .011, d = -.78, BF = 6.47). However, when including all 36 rats, all groups showed significantly more freezing to the CS+ than to the CS− (SAL: t(11) = -1.86, p = .045, d = -.33, BF = 2.01; QUIN0.05: t(11) = – 2.48, p = .015, d = -.61, BF = 4.72; QUIN5: t(11) = -2.11, p = .029, d = -.65, BF = 2.83).

During Test 3 (**Fig. 3e**), which took place in the training context (CTX-A) on Day 29, there was no evidence for Treatment x CS type interactions (QUIN0.05: F(1, 18) < .01, p = .982, η2p < .01, BF = .41; QUIN5: F(1, 16) = .74, p = .402, η2p = .04, BF = .58) and there was no statistical evidence for differences in responding to the CS+ and CS− in any of the treatment conditions (SAL: t(9) = -1.45, p = .09, d = -.5, BF = 1.25; QUIN0.05: t(9) = -1.48, p = .086, d = -.53, BF = 1.29; QUIN5: t(7) = .08, p = .531, d = .04, BF = .32).

Secondary analyses suggested no evidence for between-group differences in average freezing during CS+ presentations, baseline freezing, (i.e., before onset of the first tone), or the proportion of generalizers during any of the test sessions.

### 3.3. Experiment 4: No evidence for an effect of post-training injection of quinpirole (0.05 or 1 mg/kg) on fear generalization when using higher shock intensity during training

Experiment 4 aimed to investigate whether quinpirole injection can prevent fear generalization at a drug-free test when using a higher shock intensity during training. We expected that this higher shock intensity would result in more fear generalization at test in saline control rats than in our prior experiments (based on the results of Experiment C, see **Supplement**). Such a saline group would also be more similar to the saline condition in the mouse study by De Bundel and colleagues^13^. Note that, although at the start of our study, we hypothesized that the low dose of quinpirole (i.e., 0.05 mg/kg) would increase fear generalization, the results of Exp. 2 suggested that, if anything, this dose may prevent fear generalization. Therefore, the low dose was also used in this study, to examine if it would prevent fear generalization, similar to the hypothesis for the 1 mg/kg dose.

Despite the use of stronger shocks, saline control rats showed significantly more freezing to the CS+ than to the CS− at test (t(15) = -1.8, p = .046, d = -.35, BF = 1.78), and thus still incomplete fear generalization at the group level. Also in quinpirole rats, the difference in freezing to the CS+ and CS− was significant, and numerically even higher, especially in rats that received the dose of 1 mg/kg (QUIN0.05: t(15) = -2.34, p = .017, d = -.44, BF = 4.01; QUIN1: t(15) = -5.04, p < .001, d = -.86, BF = 410.27) (**Fig. 4a**). If significant, such increased differentiation between CS+ and CS− would be in line with our hypothesis that quinpirole should attenuate fear generalization. There was, however, no reliable evidence for Treatment (dopaminergic drug versus saline) x CS type (CS− versus CS+) interactions (QUIN (0.05 mg/kg): F(1, 30) = .25, p = .622, η2p = .01, BF = .36; QUIN (1 mg/kg): F(1, 30) = 3.28, p = .08, η2p = .1, BF = 1.08).

**Fig. 4.**
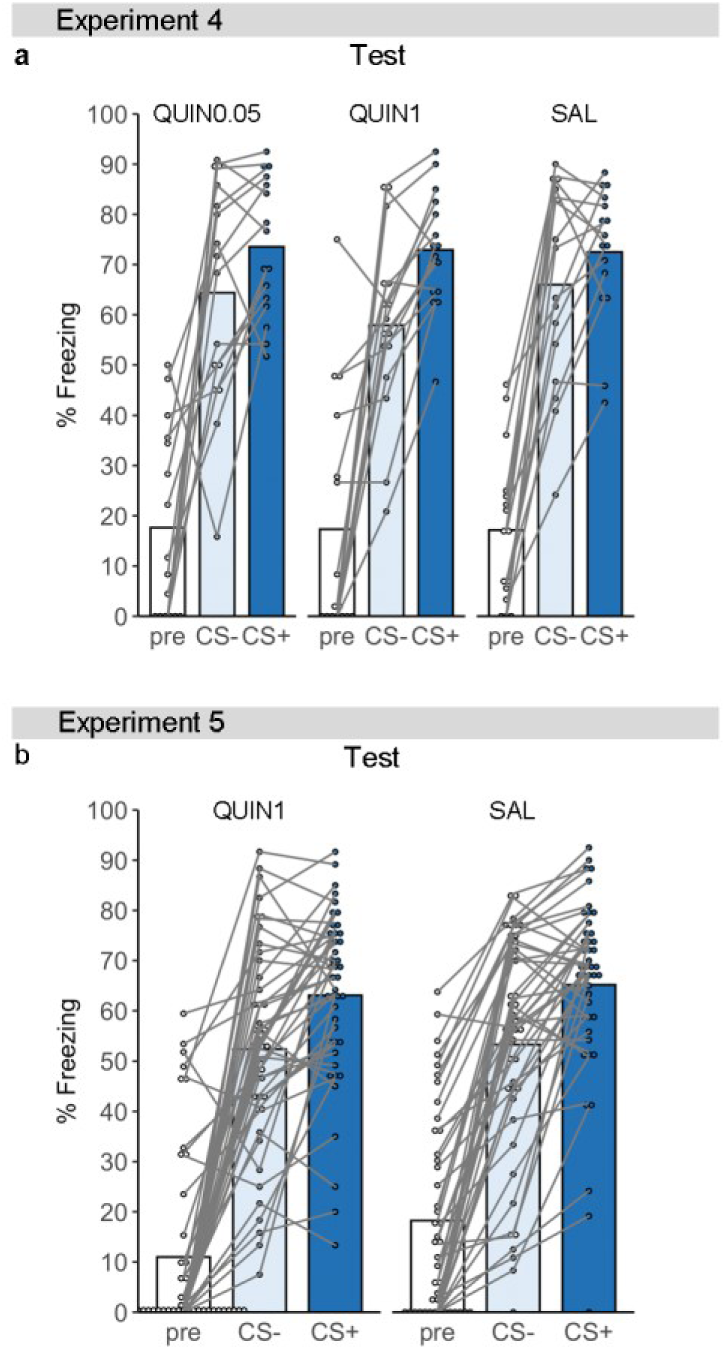
Experiments 4 & 5. Average % freezing per treatment condition during testing (Day 2; CTX-B) in Experiment 4 (panel a) and Experiment 5 (panel b), shown for the 180-s baseline period (pre), CS− and CS+. QUIN0.05 = quinpirole (0.05 mg/kg), QUIN1 = quinpirole (1 mg/kg), SAL = saline.

Secondary analyses suggested no evidence for between-group differences in average freezing during CS+ presentations, baseline freezing, (i.e., before onset of the first tone), or the proportion of generalizers.

### 3.4. Experiment 5: No evidence for an effect of post-training injection of quinpirole (1 mg/kg) on fear generalization when using higher shock intensity during training and large sample size

In Experiment 4 we observed numerically weaker generalization between the CS+ and CS− in rats that received 1 mg/kg quinpirole than in saline controls. Therefore, we aimed to investigate this effect further by taking into account the effect size observed in Exp. 4 and using a sample size that is required to detect such an effect with a power of more than 90%. Experiment 5 followed the procedures from Experiment 4, but only included two of the groups from Experiment 4 (i.e., quinpirole 1 mg/kg versus saline). The experiment had a sequential design with a predetermined maximum sample size, and was performed in three separate batches. Given that we did not observe the predefined Bayes factor for the CS type x Treatment interaction after testing batch 1 or 2, the maximum sample size of 88 rats was tested.

One saline rat was excluded from all analyses because it showed no freezing at all during testing (i.e., 0% freezing; exclusion based upon preregistered criterion). In addition, the excluded rat did not show any freezing during training either, and a lack of responding at the time the shocks should have been given, so we decided not to perform analyses with this rat included. Note that we hereby deviate from the preregistration because we planned to perform the analyses with all subjects included, apart from applying the preregistered exclusion criterion. A total of 87 rats was thus included in the analyses described below.

There was no evidence for a CS type x Treatment interaction (F(1, 85) = .17, p = .68, η2p < .01, BF = .23), and the obtained Bayes factor suggested substantial evidence for the absence of an interaction^33^ (**Fig. 4b**). One-sided paired t-tests suggested significantly more freezing to the CS+ than the CS− in saline control rats (t(42) = -4.93, p < .001, d = -.62, BF = 3032.43) and quinpirole rats (t(43) = -4.31, p < .001, d = -0.56, BF = 502.73).

Secondary analyses suggested no difference in the proportion of generalizers between saline and quinpirole rats (X^2^(1) < .01, p = 1, BF = .27). In addition, the generalization index was not significantly lower in quinpirole rats than in saline rats (t(85) = .76, p = .777, d = .16, BF = .14).

As mentioned before, we carried out an additional study (Experiment 6) to assess the acute effects of quinpirole and raclopride on locomotor activity. All details can be found in the **Supplement**. In brief, we observed locomotor effects that were largely in line with prior research. Importantly, as described in the literature ^32^, we found that 1 mg/kg quinpirole first suppressed locomotor activity, followed by a statistically significant increase in locomotor activity compared to saline (**Supplement Fig. S4**). This indicates, that in a different task than the fear conditioning procedure, the drug did produce the expected acute behavioral effect at the applied dose in male Wistar rats.

## 4. Discussion

In a series of five experiments, the dopamine D2 receptor (D2R) agonist quinpirole or D2R antagonist raclopride was injected systemically after differential cued fear conditioning in rats. One day after treatment, fear generalization was assessed by comparing average % freezing during CS+ versus CS− presentations in a new context. Our studies aimed to evaluate in rats the role of dopaminergic signaling in fear memory processing as established in mice, emulating the protocol of a previously published study by De Bundel and colleagues^13^. They showed that post-training systemic injection in mice of quinpirole (1 mg/kg) or raclopride (0.3 mg/kg) differentially affected fear responding to a learned danger cue (CS+) versus safety cue (CS−). Based on those published results, we expected quinpirole (1 or 5 mg/kg) to prevent fear generalization, and raclopride (0.3 mg/kg) to augment fear generalization. In addition, we initially hypothesized that a low dose of quinpirole (0.05 mg/kg) would also augment fear generalization, due to the assumed activation of presynaptic autoreceptors. We found no evidence in rats for an effect of systemically administered quinpirole or raclopride on fear generalization at test. In addition, there were no effects of treatment on freezing levels during CS+ presentations per se or during the baseline period at test (i.e., before tone onset) in any of the 5 experiments. We did, however, replicate previously described acute behavioral effects of quinpirole (0.05, 1 and 5 mg/kg) and raclopride (0.3 mg/kg) during a locomotor activity test, indicating that the applied drug doses and route of administration were capable of inducing previously reported behavioral effects.

In the light of the inconsistent results between our studies (in rats) and the published studies by De Bundel and colleagues (in mice)^13^, a number of differences between these two studies should be acknowledged. First and foremost is of course the difference in species, given the reported species-dependent variations in ventral tegmental area dopamine signaling^34^. The fact that DA neurons in mice and rats show different sensitivity to D2-mediated inhibition could have important implications for presynaptic or postsynaptic effects of D2R modulation. Similarly, differences in DA agonist-induced locomotor effects, prepulse inhibition, or yawning between rats and mice have been described^35–37^. Another relevant question is whether mice and rats express fear in the same manner (e.g., comparable levels of freezing versus escape behavior) during presentations of tones with different frequencies. If existent, between-species differences in prototypical fear behavior may obviously complicate the comparison of studies in rats versus mice when freezing responses to the CS+ versus CS− are used as a measure of generalization.

Furthermore, and as expected, pilot studies in our lab indicated the need for different behavioral parameters than those used in the mouse studies^13^ in order to allow for the study of fear generalization in rats (e.g., tones of 5 and 10 kHz rather than 2.5 and 7.5 kHz, shocks of .5 s rather than 2 s, and intermixed rather than blocked testing). In addition, we chose to leave out the habituation session prior to conditioning, and used a semi-random rather than an alternating order of CS+ and CS− presentations during learning. Finally, our saline control animals showed, on average, a different behavioral pattern than those in the De Bundel study. The control rats in most of our dopamine studies (except for Experiment 2) showed statistically significant higher freezing during CS+ presentations than during CS−, whereas the control mice in the prior quinpirole study did not freeze more to the CS+ than to the CS− (see Fig. 1C of De Bundel et al.^13^). Nevertheless, it is clear that our control rats showed fear generalization, as freezing to the CS− was considerable, thus fulfilling the prerequisite to observe any effect on fear generalization in the quinpirole groups. The observed variability in freezing behavior of the control animals between some of our experiments could be perceived as a limitation, but is probably a logical consequence of our attempt to tread a fine line, i.e., to create a learning experience that still contained a degree of ambiguity (with the control animals of Experiment 2 showing statistically insignificant CS+ versus CS− differentiation, and thus flipping to the other side than in Experiments A, 1 and 3, where the difference was significant). As mentioned before, we did not want to overtrain the animals in their discrimination between the CS+ and CS− (which could, for example, have been achieved with multiple training days^38^), nor did we want them to have inadequate learning opportunities to differentiate between the CS+ and CS− (in such scenario, full generalization between the CS+ and CS− might just reflect a perceptual discrimination deficit). Overall, the observed behavioral variability between experiments is most likely not a major issue, given that each of our experiments had its own appropriate saline control group.

A potential limitation of our study is that we focused on systemic drug injection and did not directly target a particular brain region via, for example, drug infusion or optogenetic manipulation. Given the apparent lack of generalizability to rats of the results of De Bundel et al.^13^ of systemic application of dopaminergic drugs, a follow-up study in rats may want to use intracerebral drug administration (as was done in mice in Fig. 2I and 3I of De Bundel et al.^13^). Such a study will allow for even stronger conclusions about the replicability (and generalizability) of the effects described in mice^13^ specifically, and about the role of post-training D2R signaling in subsequent fear generalization in rats more generally.

In summary, contrary to our hypothesis, we did not find systemic raclopride (0.3 mg/kg) to augment fear generalization, although it should be noted that this conclusion merely relies on a single study with a sample size of 12 rats per treatment condition, yielding only anecdotal evidence for the absence of a CS type x Treatment interaction effect. In contrast, we performed five studies with quinpirole, in which we used three different doses of the drug (0.05, 1, or 5 mg/kg), two different shock intensities (.4 or 1 mA), and including a final study with a large sample size yielding a power of over 90% to detect an effect if any. Moreover, Experiment 3 (N = 36) was a replication of Experiment 2 (N = 36) (except for additional tests), and Experiment 5 (N = 88) was a replication of Experiment 4 (N = 48) (except for the omission of the 0.05-mg/kg quinpirole group in Exp. 5). The final experiment provided substantial evidence (Bayesian analyses) for the absence of an effect of quinpirole (1 mg/kg) on fear responding to the CS+ versus CS−. In other words, we have gathered evidence against a preventative (or augmentative) effect of quinpirole on fear generalization under the applied conditions in rats.

## Acknowledgements

We acknowledge the financial support of the European Research Council (ERC Consolidator Grant to T. Beckers, grant number 648176), the Research Foundation – Flanders, Belgium (FWO Doctoral Fellowship to N. Schroyens, grant number 1114018N) and KU Leuven Research Grant C16/19/002 (to T. Beckers and L. Luyten). A previous version of this paper was published on bioRxiv (https://www.biorxiv.org/content/10.1101/2022.02.02.478781).

## Author contributions

Conceptualization: N.S., T.B. and L.L.; Data curation: N.S.; Formal analysis: N.S.; Funding acquisition: N.S., T.B. and L.L.; Investigation: N.S., L.V., B.Ö. and V.A.O.S.; Methodology: N.S., J.Z., D.D., T.B. and L.L.; Project administration: T.B. and L.L.; Resources: T.B.; Supervision: T.B. and L.L.; Validation: N.S.; Visualization: N.S. and L.L.; Writing – original draft: N.S. and L.L.; Writing – review & editing: N.S., L.V., B.Ö., V.A.O.S., J.Z., D.D., T.B. and L.L.

## Supplementary Methods & Results

This Supplement contains details about the fear conditioning equipment and procedure. It also contains information related to three behavioral protocol optimization studies (i.e., Experiments A-C) which were carried out before the five psychopharmacological studies in the main text (i.e., Experiments 1-5), as well as details about secondary statistical analyses for those experiments. Finally, it contains information related to a locomotor activity study that assessed acute effects of quinpirole and raclopride (i.e., Experiment 6).

## Supplementary Methods

### Experiments A-B-C & Experiments 1-5

#### Apparatus

Four chambers (Contextual NIR Video Fear Conditioning System for Rats, Med Associates Inc., St. Albans, VT, USA, 30 cm (L) x 25 cm (W) x 21 cm (H)) illuminated by infrared and white light (45 lux), and with built-in ventilation fans (±67 dB) were used. Fear conditioning took place in context A (CTX-A), consisting of a grid floor (19 rods of 4.8 mm diameter, 16 mm center to center) and a triangular-shaped black insert. Before and/or after each behavioral session, the context was cleaned and scented with diluted cleaning product (5.55 % in water). The first test session took place in context B, consisting of a white plastic floor, a white plastic curved back wall, illuminated with infrared light only, and cleaned and scented with a different cleaning product.

#### Differential cued fear conditioning

Rats were placed in context A (CTX-A) and, after a baseline period of 120 s, two tones (5 and 10 kHz, counterbalanced) were presented five times each in semi-random order (80 dB, 30 s, ITI = 60 – 160 s, order of tone presentations = CS+, CS−, CS−, CS+, CS−, CS+, CS−, CS−, CS+, CS+). In Experiments B, C, 4, and 5, the rise time of the tones was changed from .1 s to 1s to prevent interference with freezing behavior due to startle responding at tone onset. One tone (i.e., CS+) always co-terminated with a foot shock (.5 s, .4, .8 or 1 mA, see **Tables 1 & S1** for shock intensities), while the other tone (i.e., CS−) was never paired with shock. Sixty seconds after the last tone, rats were returned to their home cage. Depending on the experiment, rats then received an intraperitoneal injection of saline, quinpirole (0.05, 1 or 5 mg/kg) or raclopride (0.3 mg/kg) in 1 ml/kg (see **Table 1**).

#### Test of fear responding to the CS+ and CS−

Twenty-four hours after training, rats underwent a drug-free test in a novel context (CTX-B). After a baseline period of 180 s, the CS+ and CS− tones were repeatedly presented in a semi-random and counterbalanced order (4 presentations of each tone, 80 dB, 30 s, ITI = 80 – 140 s, see **Tables 1 & S1** for the order of tone presentations). Thirty seconds after the last tone, rats were returned to their home cage.

Depending on the experiment, additional drug-free test sessions were performed (see **Tables 1 & S1**). Two (Exp. 1) or 26 days (Exp. 2) after training, rats were re-exposed to the training context (CTX-A, Test 2), and, after a baseline period of 180 s, the CS+ and CS− tones were repeatedly presented in a semi-random and counterbalanced order (4 presentations of each tone, 80 dB, 30 s, ITI = 80 – 140 s). Thirty seconds after the last tone, rats were returned to their home cage. In Experiment 3, a second test took place in CTX-B at day 28, and a third test in CTX-A on Day 29. The order of the tones and ITIs differed between test sessions.

### Experiments A-B-C: Behavioral protocol optimization

#### Preregistration

All study protocols and statistical analyses were preregistered on the Open Science Framework (OSF) at https://osf.io/bm76c/registrations.

#### Data and code availability

All relevant datasets, analysis scripts, and results of all preregistered statistical analyses can be found on OSF at https://osf.io/bm76c.

#### Behavioral protocol optimization studies (Experiments A-C)

We performed experiments A, B and C to obtain appropriate behavioral protocols to investigate the effects of post-training dopaminergic drugs on fear memory generalization (see **Table S1** for an overview of study characteristics).

Experiment B investigated the effect of presenting CS+ and CS− tones in blocks (as was used in the study in mice) versus intermixed order at test on fear responding to both tones, and used a higher shock intensity during training compared to Exp. A (.8 mA instead of .4 mA). Experiment C investigated the effect of shock intensity (.4 versus 1 mA) on fear responding at test to CS+ and CS− tones. Smaller pilot studies were conducted to determine the optimal frequency and intensity of the tones; they are not reported here.

All rats received an intraperitoneal injection of saline (1 ml/kg) immediately after fear conditioning. The subjects, apparatus and behavioral procedures used in these studies are identical to those described above, except for the details that are listed below.

In Experiment A, a baseline period (i.e., before onset of the first tone) of 1 min. was used. This baseline period was prolonged to 3 min. in all subsequent studies in order to properly measure baseline freezing. Experiment B compared two different test sessions, i.e., with intermixed versus blocked tone presentations. The intermixed test session was as described above in the section ‘Test of fear responding to the CS+ and CS−’. In the blocked testing group, either the CS+ or the CS− was presented three times (baseline period of 180 s, 80 dB, 30 s, ITI = 80 – 140 s) during a first test session one day after training. Thirty seconds after the last tone, rats were returned to their home cage. The next day, the other tone was tested under the same conditions. The order of tone presentations (i.e., CS+ or CS− as the first tone during test) was counterbalanced.

The behavioral protocol of Exp. A was used in Experiments 1-3 (with the addition of extra behavioral testing). We next decided to switch to the behavioral protocol of Exp. C for Experiments 4-5 because, in light of the previous results, we deemed it important to investigate the effects of quinpirole with a protocol that elicits substantial fear generalization in control rats.

#### Statistical analyses

Detailed information regarding all statistical analyses can be found in the main text. Here, we only describe specifics for behavioral protocol optimization of Experiments A-C.

For Experiments B-C, rats showing less than 10% freezing on average during CS+ presentations at Test 1 were excluded from the analysis according to a preregistered exclusion criterion. For Experiments A-C, one-sided paired t-tests assessed whether freezing during CS+ was higher compared to CS− for each condition (i.e., classification into conditions based on testing intermixed versus blocked in Exp. B and shock intensity in Exp. C). In order to investigate whether shock intensity during training affected fear responding to the CS+ and CS− at test in Exp. C, we performed a mixed ANOVA with between-subjects factor Shock intensity (.4 or 1 mA) and within-subjects factor Trial type (CS− or CS+).

**Table S1.**
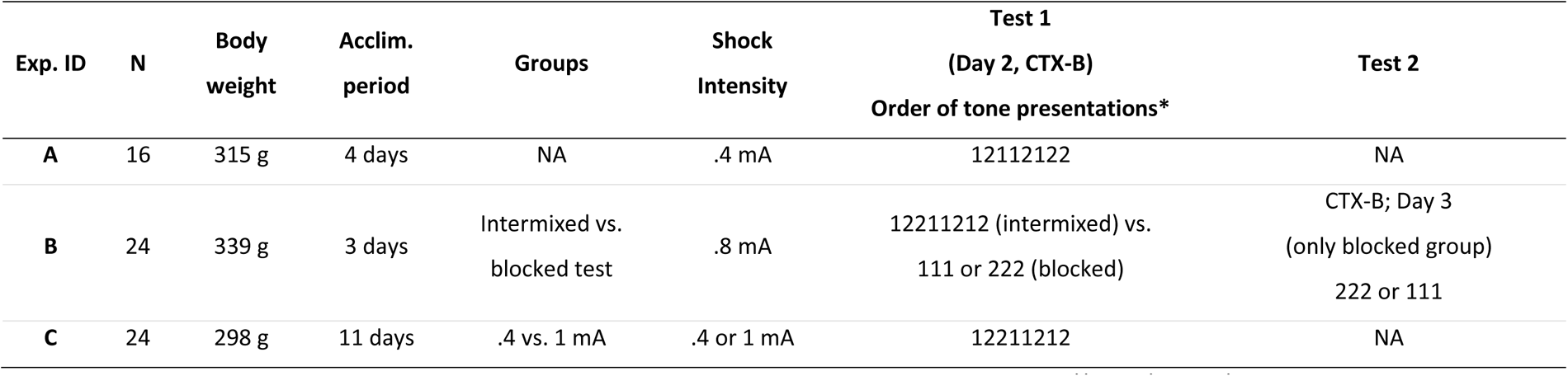
Behavioral protocol optimization studies. All experiments were preregistered on OSF at https://osf.io/bm76c/registrations. Sample size (N) and average body weight one day before training are shown. Acclim. period = interval between the arrival of the male Wistar rats in our animal facility and the start of handling (2 subsequent days of handling preceded the fear conditioning protocol). Rats were trained in context A (CTX-A) and tested in context B (CTX-B). **\***The order of the tones at test was counterbalanced (1 = CS+, 2 = CS− or 1 = CS−, 2 = CS+). During intermixed tests, 4 CS+ and 4 CS− presentations occurred in intermixed and semi-random order within one session, whereas blocked testing occurred in two separate test sessions with either 3 CS− or 3 CS+ presentations.

### Experiments 1-5: Effects of dopaminergic drugs on subsequent fear generalization

#### Statistical analyses

In addition to the primary statistical analyses described in the main text, we also conducted secondary analyses. These secondary analyses included two-sided t-tests to assess whether treatment affected average freezing (1) during CS+ presentations and (2) during baseline at test.

For Experiments 1-4, Chi-squared tests were performed to investigate whether treatment influenced the proportion of (complete) generalizers. We preregistered to calculate a generalization index (GENINDEX = average % freezing during CS− divided by average % freezing during CS+ at test) for each rat. Rats were subdivided in “differentiators” (GENINDEX smaller or equal to 0.75) and “generalizers” (GENINDEX larger than 0.75, but smaller than 1.30). Note that differentiators may actually still show partial fear generalization (if they freeze considerably to the CS−), yet this generalization is less complete than in the generalizers group. Rats showing considerably higher freezing during CS− compared to CS+ (GENINDEX larger or equal to 1.30) were not included in the Chi-Squared tests, as they could not be unambiguously classified as generalizers or differentiators.

Secondary analyses for Experiment 5 included calculation of the generalization index (GENINDEX = average % freezing during CS− divided by average % freezing during CS+) for each rat, and rats were subdivided into “differentiators” (GENINDEX smaller or equal to 0.90) and “generalizers” (GENINDEX larger than 0.90, but smaller than 1.10). Rats showing considerably higher freezing during CS− compared to CS+ (GENINDEX larger or equal to 1.10) were not included in the Chi-Squared tests, as they could not be unambiguously classified as generalizers or differentiators. The rationale for changing the criteria for classification into generalizers and differentiators can be found in the preregistration form for Experiment 5 at https://osf.io/gnqpw. In brief, it was revised because of the higher shock intensity and expected higher overall freezing values (as based upon Exp. C and 4) in Exp. 5 in comparison with earlier experiments (Exp. 1-3). As in our previous studies, Chi-Squared tests (quinpirole versus saline) were performed to assess whether quinpirole treatment affected the proportion of generalizers. In addition, we conducted one-sided t-tests to investigate whether the GENINDEX was lower in quinpirole-treated rats than in saline control rats.

### Experiment 6: Acute effects of raclopride and quinpirole on locomotor activity

#### Preregistration

All study protocols and statistical analyses were preregistered on the Open Science Framework (OSF) at https://osf.io/9d72h/registrations.

#### Data and code availability

All relevant datasets, analysis scripts, and results of all preregistered statistical analyses can be found on OSF at https://osf.io/9d72h.

#### Rationale for the evaluation of the acute effects of raclopride and quinpirole on locomotor activity

Previous research in rats has shown acute effects of systemic administration of raclopride and quinpirole on locomotor activity^1–6^. As a positive control of drug efficacy, non-naïve rats received an intraperitoneal saline or drug injection and acute effects on locomotor activity were assessed (Experiment 6). Based on the previously mentioned literature, we hypothesized that raclopride would generally suppress locomotor activity during a 1-h test session. Regarding quinpirole, it has been suggested that a dose of 1 mg/kg would first activate presynaptic autoreceptors, resulting in a transient decrease in dopamine synthesis and locomotor activity, followed by activation of postsynaptic dopamine D2 receptors, inducing hyperactivity^5^. On the other hand, a low dose of quinpirole (0.05 mg/kg) would solely activate presynaptic autoreceptors. Thus, quinpirole was expected to generally suppress locomotor activity at a dose of 0.05 mg/kg, whereas higher doses (1 or 5 mg/kg) would first suppress and later increase locomotor activity.

#### Subjects

At the age of 15-16 weeks, 35 rats that previously underwent a fear conditioning procedure (identical to the one adopted in Experiment 2) were used to test the acute drug effects during a locomotor activity test.

#### Drug administration

Rats received an intraperitoneal injection of saline, 0.3 mg/kg raclopride, 0.05 mg/kg quinpirole, 1 mg/kg quinpirole, or 5 mg/kg quinpirole (1 ml/kg; 7 rats/treatment condition).

#### Apparatus

Five minutes after the injection, rats were exposed to a novel testing box (38 x 38 x 60 (h) cm) for 1 h to assess effects on locomotor activity (distance travelled and % movement). Rat behavior was recorded by a video camera that was mounted above the experimental set-up. Testing occurred in the dark. Distance travelled (cm) and % movement during the 1h-session were measured in 5-min time bins using Ethovision software XT (version 11.5, Noldus, Netherlands). The software’s default criteria, i.e., start velocity of 2 cm/s and stop velocity of 1.75 cm/s, were used to calculate % movement.

#### Statistical analyses

Mixed Treatment x Time (twelve 5-min time bins) ANOVAs were used to assess treatment effects over time for raclopride (RACLO, SAL) and quinpirole (SAL, QUIN0.05, QUIN1, QUIN5) separately. Significant ANOVAs were followed up by Tukey post hoc tests to assess effects of treatment during each 5-min time bin, except for the first and the last time bin given that we had registered planned comparisons (one-sided t-tests) at those time points based on our a priori hypotheses (see https://osf.io/5bsh7). Specifically, we performed one-sided t-tests to assess treatment effects during the first and last 5 min of the test session. For the first 5 min (i.e., min 0-5) we expected for distance travelled and % movement a general suppression of locomotor activity by all treatment conditions, compared to saline (i.e., SAL > RACLO, QUIN0.05, QUIN1, QUIN5) and for the last 5 min (i.e., min 55-60) we expected an enhancement of locomotor activity in rats that received 1 and 5 mg/kg quinpirole, compared to saline (i.e., SAL < QUIN1, QUIN5).

## Supplementary Results

### Experiments A-B-C: Behavioral protocol optimization

#### Experiment A: Inducing intermediate levels of fear generalization

The aim of Experiment A was to obtain a behavioral protocol that could be used to investigate the effects of post-training dopamine D2 receptor (ant)agonism on later fear responding to the CS+ and CS−. Rats were trained using a differential cued fear conditioning protocol, injected intraperitoneally with saline immediately after training, and tested one day later in a new context B (CTX-B) to assess the amount of fear generalization between CS+ and CS−. About half of the rats showed higher freezing during the CS+ than the CS− at test, whereas the other half showed (almost) complete fear generalization (i.e., no or negligible difference in freezing between CS+ and CS−, see **Fig. S1**). Average freezing was significantly higher during CS+ presentations, compared to CS− presentations (one-sided paired t-test; t(15) = -2.08, p = .028, d = -.33, BF = 2.67), indicating that, at the group level, generalization was not complete. We decided to use this protocol in Experiments 1, 2, and 3 as there were no apparent ceiling or floor effects, thus theoretically allowing for the bidirectional modulation of generalization (i.e., an increase or decrease in freezing and/or generalization resulting from the drug injection after training).

**Figure S1.**
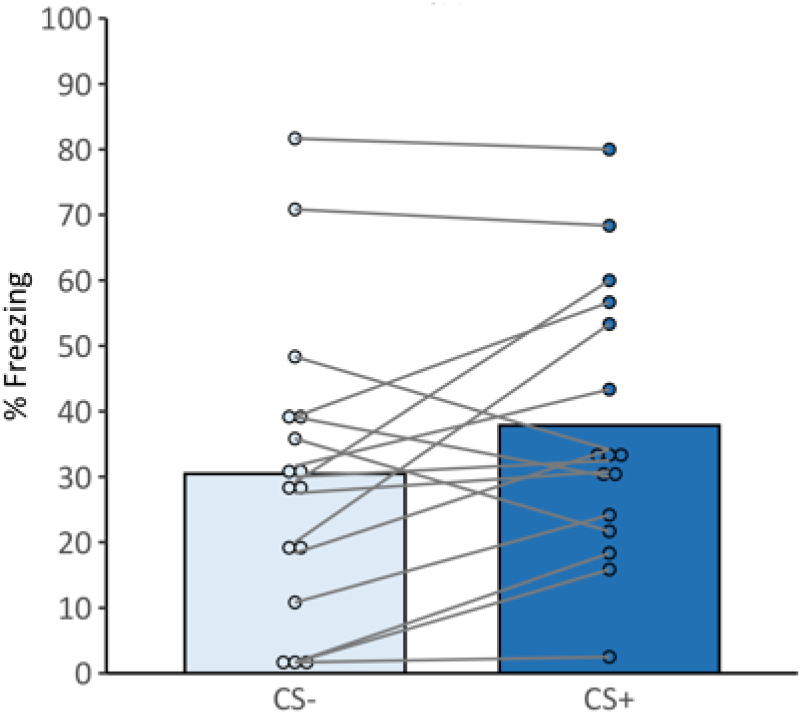
Average % freezing during testing (Day 2; CTX-B) of Experiment A.

#### Experiments B and C: Inducing stronger fear generalization

The aim of Experiments B and C was to obtain a behavioral protocol that would induce stronger generalization between CS+ and CS− at test than in Experiment A, i.e., no statistically significant difference in freezing levels during CS+ versus CS− at the group level. Such a behavioral pattern would be more in line with what was observed in mice in the study by De Bundel et al.^7^, who reported similar freezing levels during the CS+ and CS− in saline control animals (and prevented this maximal level of fear generalization by post-training quinpirole injection; see Fig. 1C by De Bundel et al, 2016). To this aim, we increased shock intensity during training (i.e., from .4 mA used in Exp. A (as well as Exp. 1-3) to .8 mA in Exp. B and 1 mA in Exp. C (as well as Exp. 4-5)). In addition, Experiment B aimed to compare freezing levels between two different types of test sessions. Intermixed tone presentations (i.e., CS+ and CS− presented in intermixed and semi-random order, in a single session) were compared to blocked tone presentations (i.e., CS+ and CS− presented in separate test session on different days, similar to De Bundel et al.^7^).

The results of Experiment B (**Fig. S2**) show that, despite the use of a higher shock intensity, rats still showed significantly higher freezing during CS+ than CS− (one-sided t-tests; intermixed test session: t(11) = -2.62, d = -.48, p = .012, BF = 5.82; blocked test: t(11) = -2.67, d = -.59, p = .011, BF = 6.24). A visual analysis suggested that the freezing levels observed during the blocked and intermixed tests were similar. Exploratory analyses suggested that there was a significant effect of testing order (i.e., whether CS+ or CS− was presented as the first tone during testing) on the amount of fear generalization when using the blocked tests, as suggested by a significant Testing order (CS− first versus CS+ first) x CS type (CS− versus CS+) interaction (F(1, 68) = 6.50, p = .013, η2p = .087, BF = 3.94). Visual analysis of supplemental graphs (see https://osf.io/kg8vb/, section 5.3) illustrates that, on average, rats that received CS+ presentations during the first test show a larger difference in responding to the CS+ and CS−, than rats that received CS− presentations during the first test. There was no evidence for such an effect of testing order on generalization when using the intermixed test (F(1,92) = .42, p = .521, η2p = .004, BF = .36).

**Figure S2.**
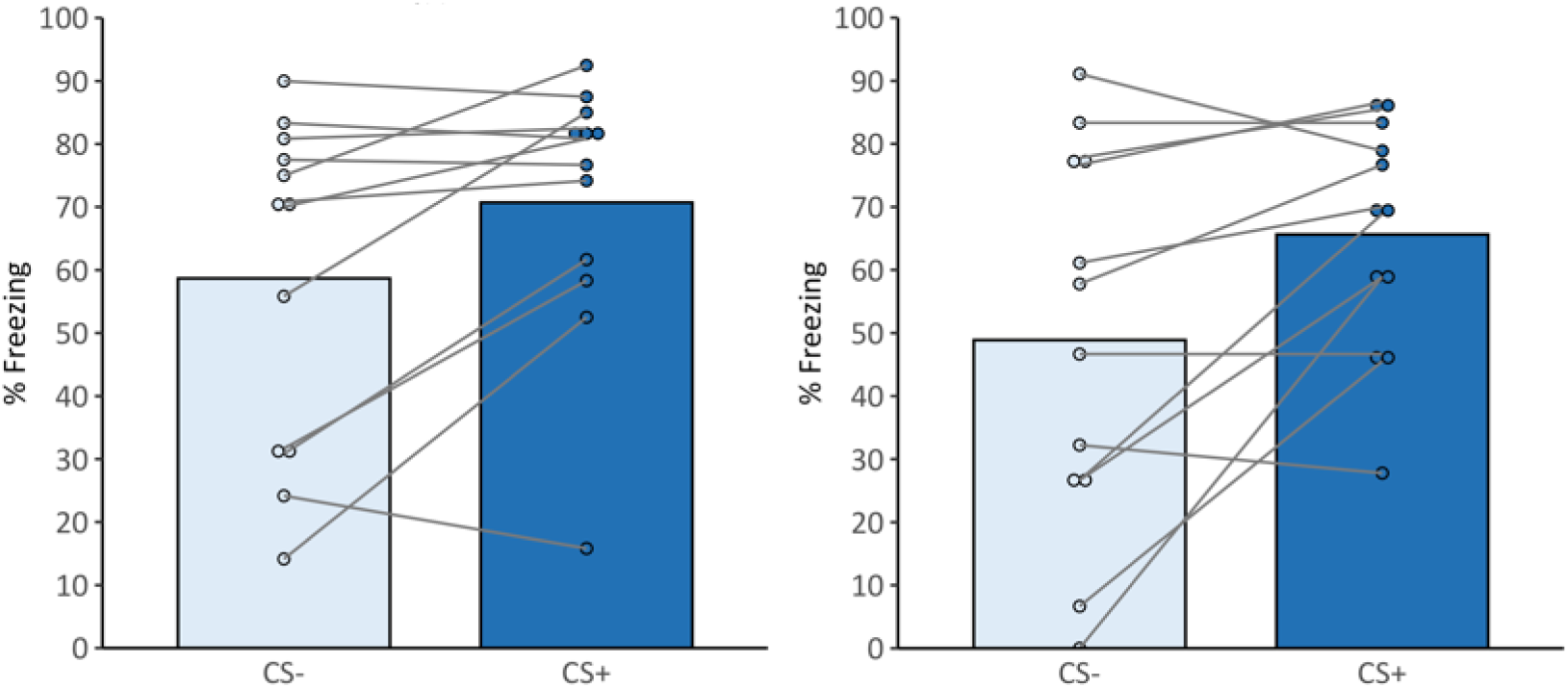
Average % freezing during intermixed testing (Day 2; CTX-B; left panel) and blocked testing (Day 2 and 3; CTX-B; right panel) of Experiment B.

In Experiment C (**Fig. S3**), two different shock intensities were used during training, i.e., .4 mA and 1 mA. There was a statistically significant difference in freezing between CS+ and CS− presentations at test if shocks of .4 mA were used during training, in line with what we observed in Experiment A (one-sided t-test; t(11) = -7.11, d = -1.1, p < .001, BF = 2380.46), whereas there was no statistically significant difference in freezing between CS+ and CS− when 1-mA shocks were used (one-sided t-test; t(11) = -1.23, d = -.34, p = .122, BF = 0.92). In addition, there was a significant Shock intensity x CS type interaction (F(1,22) = 20.37, η2p = .48, p < .001, BF = 394.35). To conclude, increasing the shock intensity during training from .4 to 1 mA significantly increased generalization at test. The behavioral protocol from Experiment C (with 1-mA shocks) was therefore used in Experiments 4 and 5.

**Figure S3.**
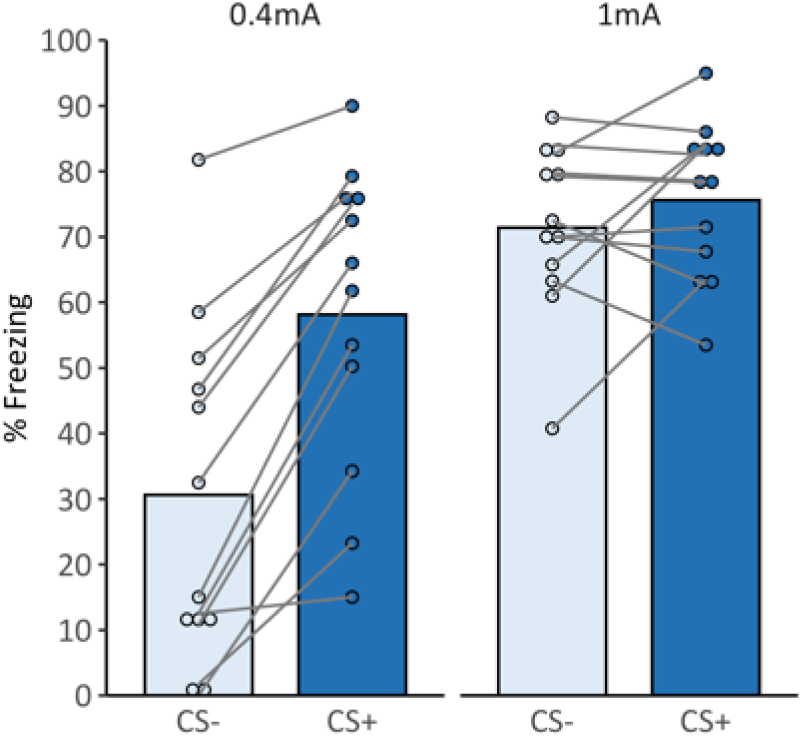
Average % freezing scores during testing (Day 2; CTX-B) of Experiment C illustrate that rats receiving 1-mA shocks during training show more generalization between CS+ and CS− than rats that received .4-mA shocks.

### Experiment 6: Acute effects of raclopride and quinpirole on locomotor activity

#### Raclopride (0.3 mg/kg) and quinpirole (0.05 mg/kg) temporarily suppress locomotor activity

In line with previously published studies and our preregistered hypotheses (see Methods section), raclopride (0.3 mg/kg) and the low dose of quinpirole (0.05 mg/kg) induced suppression of locomotor activity (as compared to saline, **Fig. S4**). For both drugs, mixed ANOVAs showed statistically significant main effects of Treatment and Time, and a significant interaction (see **Table S2**). Tukey post-hoc tests showed that the raclopride- and quinpirole-induced differences in distance travelled and % movement compared to saline rats were temporary and became statistically insignificant during the second half of the test session. These findings corroborate previous research and illustrate that the drugs produced the expected acute behavioral effects at the applied doses in Wistar rats.

**Figure S4.**
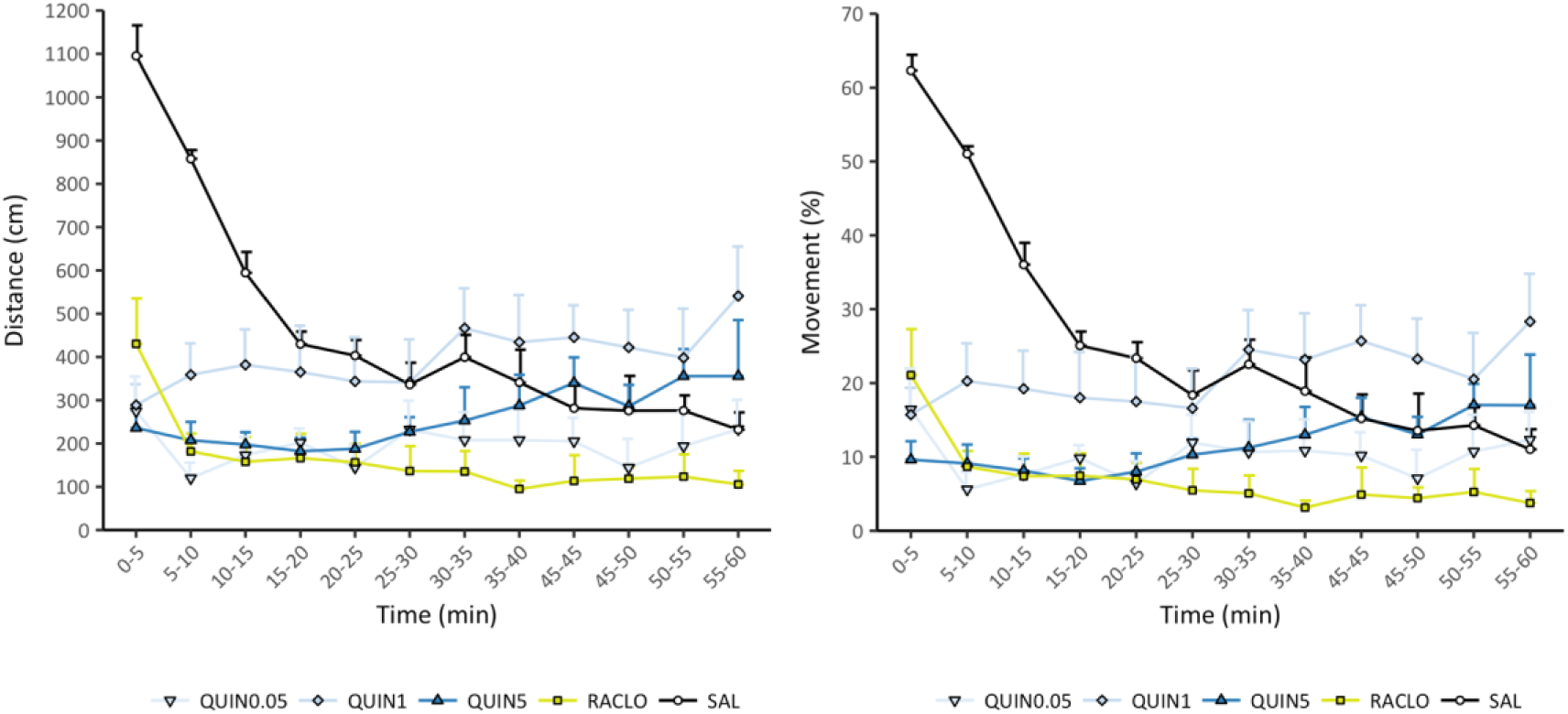
Acute effects of raclopride (RACLO; 0.3 mg/kg) and quinpirole (QUIN, different doses in mg/kg) on locomotor activity (means and SEMs) in Experiment 6. Distance travelled (cm) (left panel) and % movement (right panel) were measured to assess locomotor activity.

#### An intermediate dose of quinpirole (1 mg/kg) first suppresses and later increases locomotor activity

In line with previously published studies and our preregistered hypotheses (see Methods section), the intermediate quinpirole dose (1 mg/kg) first suppressed locomotor activity, followed by a statistically significant increase in locomotor activity compared to saline (**Fig. S4**). One-sided t-tests suggested that during the first 5 min of the test (i.e., 5-10 min after injection), quinpirole rats showed a statistically significant decrease in distance travelled and % movement compared to saline rats, whereas they showed a statistically significant increase compared to saline rats during the last 5 min of the test (see **Table S3**). This is in line with previous research and illustrates that quinpirole (1 mg/kg) produced the expected acute behavioral effects.

#### A high dose of quinpirole (5 mg/kg) temporarily suppresses locomotor activity

We assumed that the biphasic behavioral pattern that was observed after injection of an intermediate 1 mg/kg-dose of quinpirole (i.e., brief suppression followed by an increase in locomotor activity after injection) would be even more pronounced when using a higher dose (5 mg/kg). One-sided t-tests suggested that the highest dose of quinpirole induced a statistically significant suppression of locomotor activity (i.e., distance travelled and % movement) during the first 5 min. of the test session (**Fig. S4**, **Table S3**). However, contrary to our expectations, its stimulating effects at the end of the test session (i.e., about 1 hour after injection) were only moderate and insignificant when compared to saline.

**Table S2.**
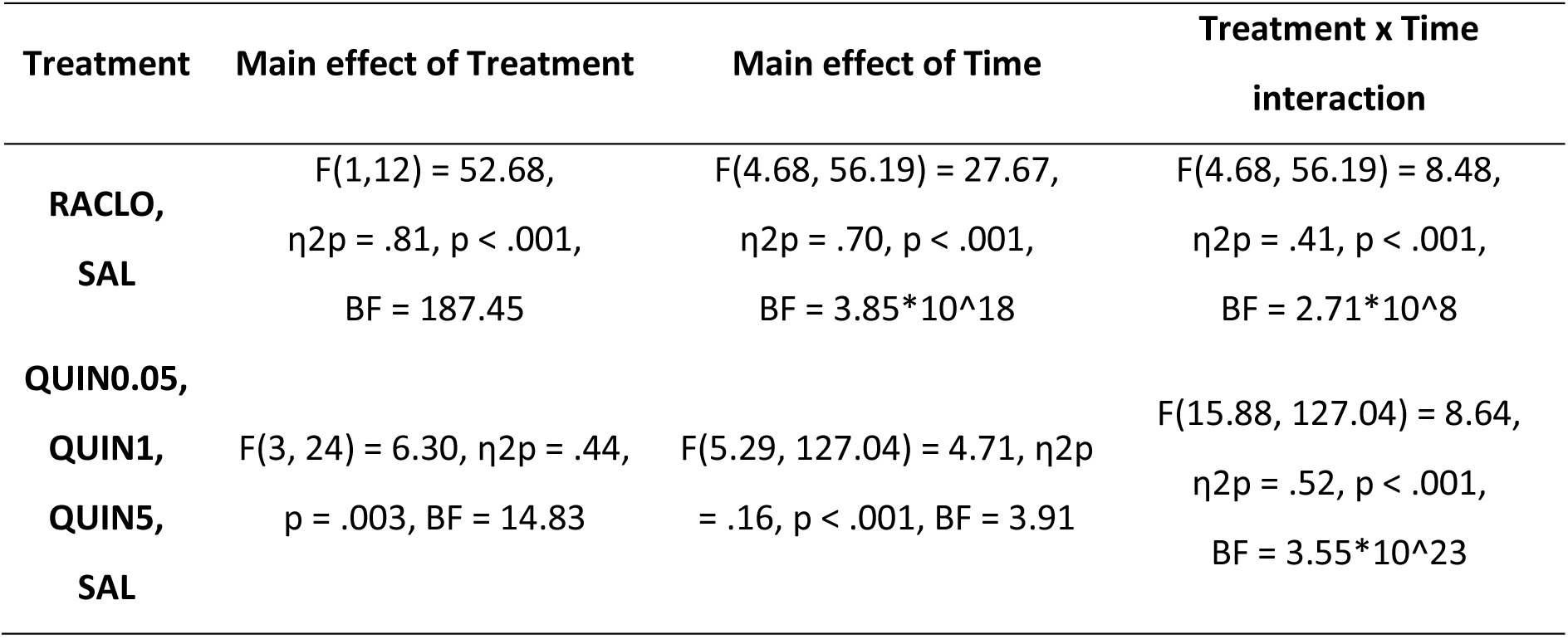
Results of preregistered mixed ANOVAs show statistically significant main effects of Treatment and Time, and significant Treatment x Time interactions for raclopride and quinpirole during the locomotor activity test (Experiment 6).

**Table S3.** Results of preregistered one-sided t-tests during the first and last five minutes of the locomotor activity test (Experiment 6).

| Treatment | First 5 min. of test session (min. 0-5) | Last 5 min. of test session (min. 55-60) |
| --- | --- | --- |
| RACLO | $t(12) = 5.26$, $d = -2.81$, $p < 0.001$,<br>BF = 103.19 | NA |
| QUIN0.05 | $t(12) = 7.66$, $d = -4.10$, $p < 0.001$,<br>BF = 2049.88 | NA |
| QUIN1 | $t(12) = 9.45$, $d = -5.05$, $p < 0.001$,<br>BF = 13802.42 | $t(12) = 2.55$, $d = 1.36$, $p = 0.013$,<br>BF = 2.76 |
| QUIN5 | $t(12) = 10.29$, $d = -5.50$, $p < 0.001$,<br>BF = 31009.8 | $t(12) = 0.91$, $d = 0.49$, $p = 0.190$,<br>BF = 0.58 |

